# rAAV2-retro-Mediated Artificial miRNA Delivery for Spinal Motor Neuron-Targeted Gene Silencing in a Neonatal ALS Mouse Model

**DOI:** 10.1101/2025.10.17.682996

**Authors:** Xueqi Gong, Yecheng Xie, Hao Wei, Qin Wang, Jiayi Wu, Zeyi Bian, Wenyuan Wang, Tonghui Xu

**Affiliations:** Laboratory Animal Center, Fudan University, Shanghai 200433, China; Laboratory Animal Resource Center, Fudan University, Shanghai 200433, China.; School of Life Sciences, Fudan University, Shanghai 200438, China; Department of Neurology, Children’s Hospital, Zhejiang, University School of Medicine, National Clinical Research, Center for Child Health, Hangzhou 310052, China; Interdisciplinary Research Center on Biology and Chemistry, Shanghai Institute of Organic Chemistry, Chinese academy of Science, Shanghai 200032, China; Department of Rehabilitation Medicine, Huashan Hospital, Fudan University, Shanghai 200040, China

**Keywords:** AAV vectors, retrograde transport, gene silencing, artificial microRNA, SOD1, spinal motor neuron, ALS mouse model

## Abstract

Amyotrophic lateral sclerosis (ALS) is a progressive and fatal neurodegenerative disorder with limited therapeutic options. A major challenge for developing effective therapies is the safe and efficient delivery of genetic cargo to the spinal motor neurons that degenerate in the disease. This is particularly pertinent for *SOD1*-linked familial ALS, where gene silencing approaches are promising but remain constrained by suboptimal delivery. Here, we establish a gene delivery platform that overcomes this barrier by combining the retrograde transport of rAAV2-retro with an artificial miRNA to achieve pan-spinal *SOD1* silencing. A single intramuscular injection of rAAV2-retro-miRNA into neonatal *SOD1*^G93A^ mice achieved widespread transduction of spinal motor neurons, significant reduction of mutant SOD1 protein, and conferred multifaceted therapeutic benefits. Treated mice exhibited delayed disease onset, extended lifespan, preserved motor function, reduced neuroinflammation, and protection of neuromuscular junctions and spinal motor neurons. These findings establish the rAAV2-retro-miRNA platform as a versatile and efficient tool for spinal motor neuron-targeted gene silencin*g,* addressing the delivery challenge and providing a foundation for further development in motor neuron diseases.

## Introduction

Amyotrophic lateral sclerosis (ALS) is a progressive and fatal neurodegenerative disorder characterized by the selective and sequential degeneration of upper motor neurons in the cerebral cortex and lower motor neurons in the brainstem and spinal cord.^1^ Despite significant research efforts, the molecular mechanisms underlying ALS remain incompletely understood. Clinically, the disease manifests with the insidious onset of muscle weakness progressing to paralysis, resulting in respiratory failure and death typically within 2-5 years of symptom onset. The multifactorial pathophysiology of ALS encompasses a complex interplay of converging mechanisms, including dysregulated RNA metabolism, aberrant protein homeostasis, impaired nucleo-cytoplasmic transport, defective axonal trafficking, mitochondrial dysfunction with concomitant oxidative stress, glutamate-mediated excitotoxicity, and chronic neuroinflammation.^2^ Current management is largely palliative, focusing on symptomatic control and ventilatory support,^3, 4^ with no therapy has been shown to halt or reverse disease progression. Pharmacological interventions such as riluzole, edaravone (Radicava), and sodium phenylbutyrate-taurursodiol modestly delay disease progression by attenuating glutamatergic transmission or mitigating oxidative stress, but their overall clinical efficacy remains limited.^4–6^

Approximately 10% of ALS cases are familial, reflecting a strong genetic contribution, with identified mutations accounting for about 70% of familial cases.^7^ Among these, mutations in genes encoding superoxide dismutase 1 (*SOD1*),^8^ TAR DNA-binding protein 43 (*TDP-43*),^9,10^ fused in sarcoma (FUS),^11, 12^ and the C9ORF72 hexanucleotide repeat expansion,^13, 14^ represent the principal molecular drivers of familial ALS. *SOD1* mutations were the first to be identified and remain among the most extensively studied genetic causes.^8, 15–17^ Recently, tofersen, an FDA-approved antisense oligonucleotide (ASO) targeting *SOD1*, achieved up to 36% reduction in the cerebrospinal fluid (CSF) SOD1 protein levels at the highest tested dose,^18, 19^ underscoring the therapeutic feasibility of gene-targeted approaches in ALS. Although its long-term clinical benefit remains to be fully established, given the limited trial cohort,^18^ this approval provides compelling validation for molecular therapies that modulate pathogenic gene expression. In this context, gene delivery has emerged as a promising approach to modulate pathogenic gene expression, including the delivery of functional genes, neuroprotective factors, or gene-silencing agents.

In the context of gene therapy for neurodegenerative disorders such as ALS, adeno-associated virus (AAV) vectors have emerged as the widely used gene delivery platforms owing to their favorable safety profile, minimal immunogenicity, and ability to sustain long-term transgene expression across cell populations within the central nervous system (CNS).^20, 21^ This inherent tropism for neural tissue enables the precise implementation of gene silencing strategies. The high transduction efficiency of AAV vectors in neural cells makes them particularly well-suited for delivering gene-silencing constructs that target mutant cytosolic *SOD1*, a well-established genetic driver of ALS pathogenesis and a central focus of therapeutic innovation.^8, 15–17^ To achieve targeted gene silencing, artificial microRNAs (miRNAs) have emerged as a promising approach owing to their superior biosafety and reduced cytotoxicity, ^22, 23^ making them particularly suitable for sustained applications in chronic neurodegenerative diseases such as ALS.

Achieving efficient and sustained transgene expression within disease-relevant compartments, particularly spinal motor neurons spanning the lumbar, thoracic, and cervical segments, remains a major challenge in ALS gene therapy. Current delivery modalities - including intracerebroventricular (ICV), intrathecal (IT), spinal subpial, and intraparenchymal injections, are constrained by significant translational barriers - require invasive neurosurgical procedures and pose considerable safety concerns. In contrast, intramuscular (IM) administration offers a peripheral route that avoids the need for neurosurgical procedures. However, prior studies delivering AAV-encoded neuroprotective factors (such as IGF1, IGF2, and HGF) via IM routes have yielded variable therapeutic efficacy in *SOD1*^G93A^ models, likely attributable to suboptimal transduction efficiency of spinal motor neurons.^24–28^ Notably, although a lentiviral-based approach has been reported to achieve SOD1 silencing in neonatal ALS mice, AAV-mediated miRNA delivery via the neonatal intramuscular route has not been previously explored or validated in ALS.

To overcome the limitations associated with current CNS delivery approaches, this study developed a gene-silencing strategy employing rAAV2-retro vectors administered via IM injection in the *SOD1*^G93A^ mouse model of ALS. Our prior work demonstrated that a single IM injection of rAAV2-retro into neonatal mice enables widespread transduction of motor neurons across the spinal cord and brainstem motor nuclei, conferring three principal advantages for motor neuron-targeted delivery, including efficient retrograde transduction of lower motor neurons, simple peripheral delivery, and intrinsic tropism for motor neurons, minimizing off-target effects.^29^ This established neonatal intramuscular injection as a well-defined system for validating the platform’s transduction and silencing efficacy in a proof-of-concept setting. Based on this delivery platform, we hypothesized that a single neonatal IM injection of rAAV2-retro encoding an artificial miRNA could achieve pan-spinal silencing of the mutant gene, thereby delaying disease onset and extending survival in *SOD1*^G93A^ mice. The *SOD1*^G93A^ model, which constitutively expresses mutant human SOD1(hSOD1), ^30^ recapitulates core ALS pathophysiology, including progressive motor neuron degeneration, paralysis, and premature death, rendering it a well-validated model to test this hypothesis. We demonstrate that this strategy achieves extensive spinal cord transduction and robust *SOD1* suppression, resulting in delayed disease onset, preserved motor function, attenuated neuropathology, and prolonged survival.

## Results

### rAAV2-Retro-Mediated miRNA Delivery Achieves Efficient Spinal Cord Transduction and *SOD1* Silencing

To evaluate the efficiency of delivering an artificial miRNA targeting human *SOD1* (h*SOD1*) to the spinal cord, we used IM injection of rAAV2-retro in *SOD1*^G93A^ mice. This approach takes advantage of the strong retrograde transport capability of rAAV2-retro, enabling effective transduction of spinal motor neurons through a peripheral route to target the CNS.^29^ An artificial miRNA was engineered on the human miR-30a scaffold, as previously described,^31^ exhibiting full complementarity to h*SOD1* mRNA and containing at least four base mismatches with mouse SOD1 to ensure species-specific silencing (Figure 1A). This miRNA expression cassette, along with an eGFP reporter gene, was cloned into the rAAV2-retro vector under the control of a CAG promoter (designated rAAV2-retro-miRNA). A control vector expressing a scrambled miRNA sequence (rAAV2-retro-Scramble) was generated in parallel.

**Figure 1.**
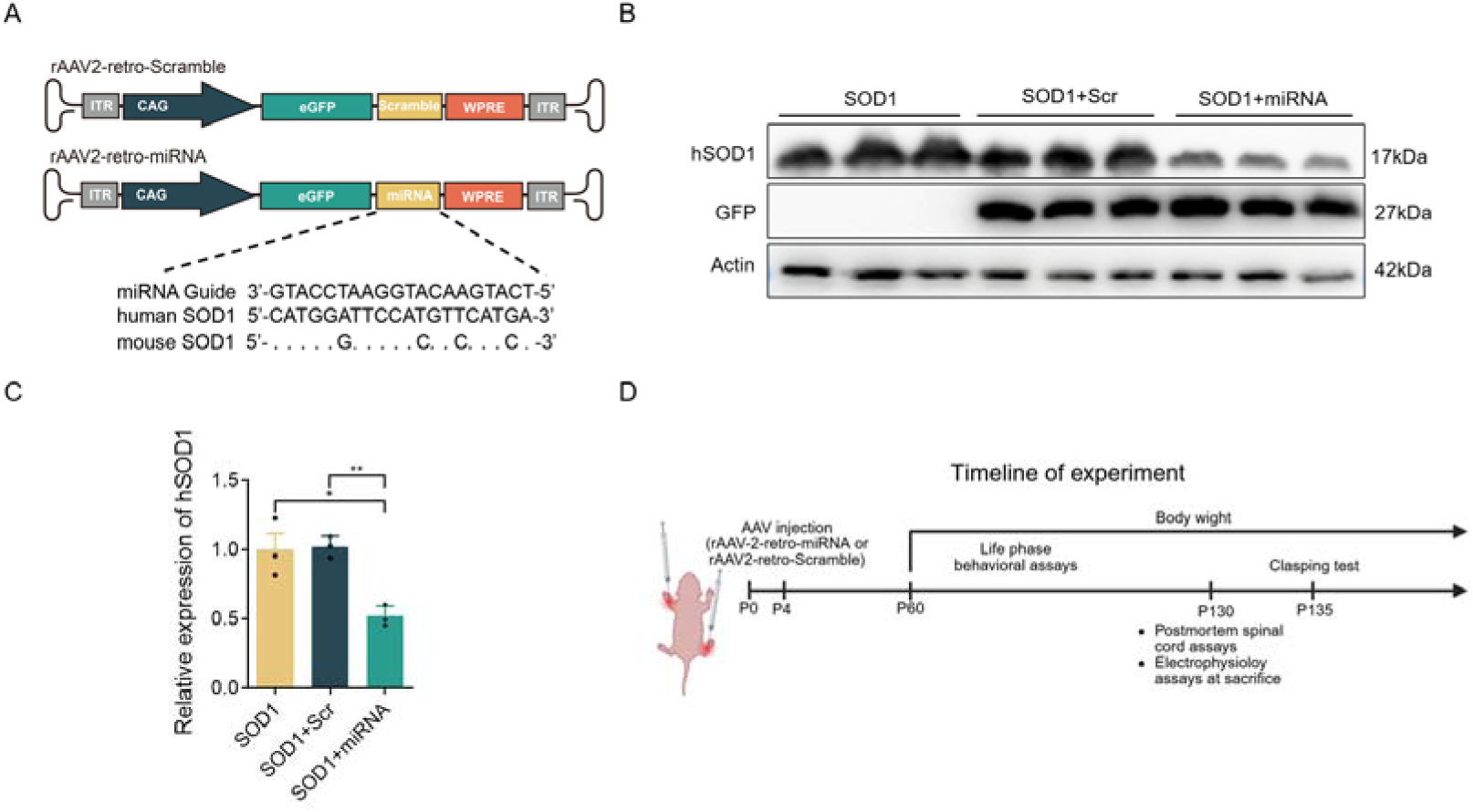
Design and *in vivo* validation of miRNA-hSOD1, an artificial miRNA targeting h*SOD1*. (A) Schematic of the AAV vector expressing miRNA-hSOD1. The artificial miRNA was designed with perfect complementarity to h*SOD1*and based on the backbone of cellular miR-30a. A scrambled miRNA that does not target any known gene product was used as a negative control. WPRE: woodchuck hepatitis post-transcriptional regulatory element; ITR: inverted terminal repeat. (B) Western blot analysis of hSOD1 protein expression in spinal cord tissues from rAAV2-retro-Scramble-treated, or rAAV2-retro-miRNA-treated ALS mice. (C) Quantification of h*SOD1* suppression in spinal cord tissues. Data are presented as means ± SEM, n=3 mice per group. One-way ANOVA with Tukey’s post hoc test. \**P* < 0.05, \*\**P* < 0.01. (D) Schematic overview of the experimental timeline and analytical approaches used for ante-mortem and post-mortem evaluations.

To evaluate the silencing efficacy, *SOD1*^G93A^ mice (n = 4) received bilateral IM co-injections of rAAV2-retro-miRNA into the extensor carpi radialis (forelimb) and gastrocnemius (hindlimb) muscles at postnatal day 4 (P4; 1 µL per site, 1.0 × 10¹³ GC/mL). Four weeks post-injection, Western blot analysis of lumbar spinal cord lysates demonstrated a significant reduction (48.2%) in h*SOD1* protein levels compared to rAAV2-retro-Scramble-treated animals (Figures 1B and 1C). The expression of endogenous mouse *Sod1* mRNA remained unchanged (Figure S1), confirming its specificity for the human transgene.

Following confirmation of target knockdown, the therapeutic potential of this strategy was assessed. *SOD1*^G93A^ mice of both sexes were treated at P4 with IM injections of either rAAV2-retro-miRNA or rAAV2-retro-Scramble (1 µL per muscle, 1.0 × 10¹³ GC/mL). Longitudinal evaluations included disease onset (defined by peak body weight decline), survival, motor performance (rotarod, grip strength, open-field tests), and neurological signs (limb clasping). Terminal electrophysiological (compound muscle action potential, CMAP) and histological analyses were conducted at 130 days of age (Figure 1D).

### Intramuscular Delivery of rAAV2-Retro-miRNA Significantly Delays Disease Onset and Extends Survival in *SOD1*^G93A^ Mice

To assess the therapeutic efficacy of rAAV2-retro-mediated *SOD1* silencing, disease progression in treated *SOD1*^G93A^ mice was longitudinally monitored by measuring body weight and survival until the humane endpoint (Figure 2 and Figure S2). Disease onset was defined as the age at which animals sustained a 10% loss from their peak body weight. Remarkably, rAAV2-retro-miRNA treatment significantly delayed disease onset by 38 days compared to scramble-injected controls (median onset: 171.5 vs. 133.5 days; *P* < 0.0001; Figure 2A). Moreover, miRNA-treated mice exhibited a pronounced extension in median survival, living 61 days longer than scramble-injected control animals (median survival: 216 days vs. 155 days; *P* < 0.0001; Figures 2B). Consistently, the mean age of death was significantly higher in the miRNA-treated group compared to the scramble controls (147.6 vs. 200.7 days; P < 0.0001; Figure 2C).

**Figure 2.**
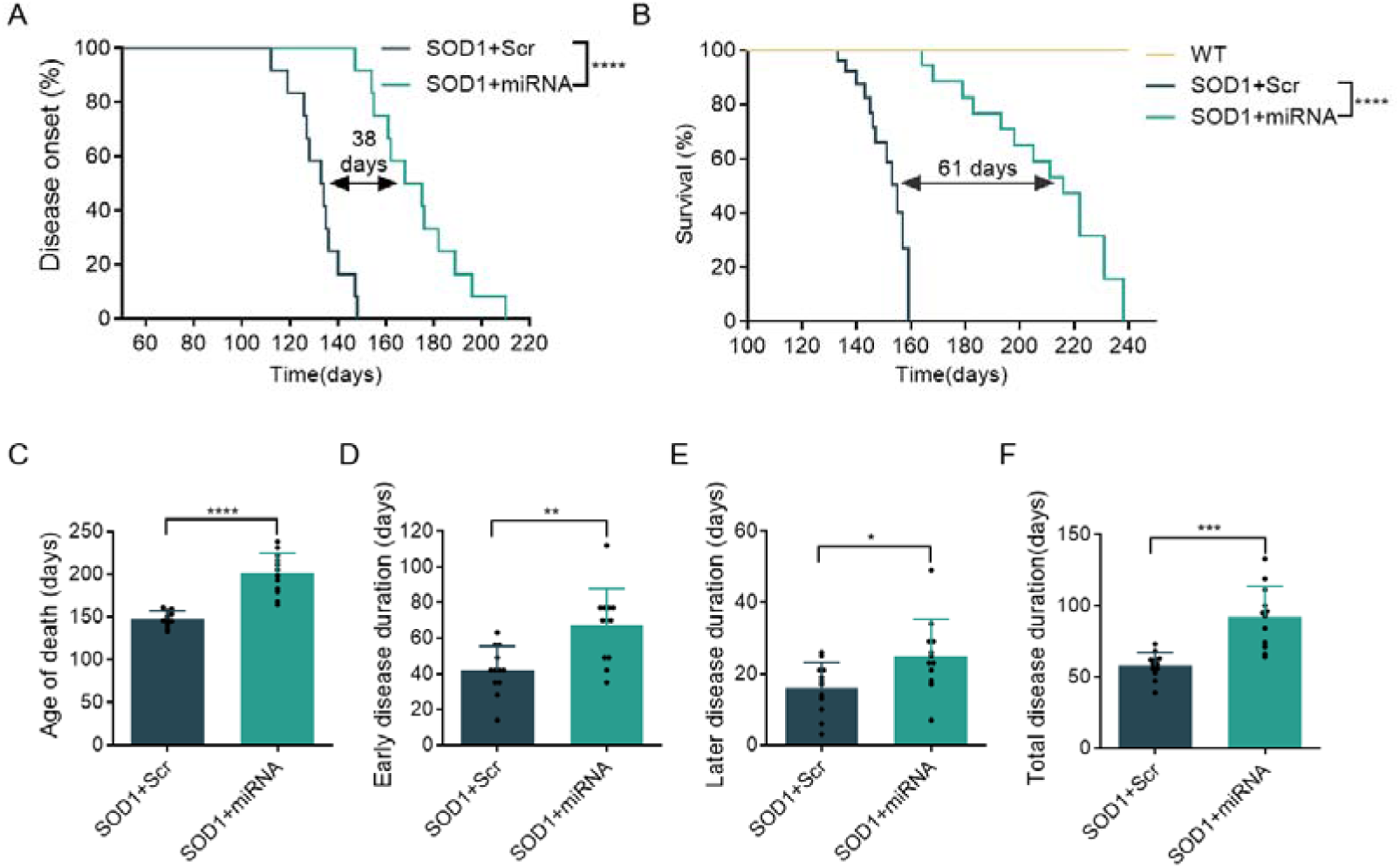
Intramuscular delivery of rAAV2-retro-miRNA delays paralysis, prolongs disease progression, and improves survival in neonatal *SOD1*^G93A^ mice. (A) Kaplan-Meier curves showing disease onset, defined as a 10% decrease from peak body weight, in rAAV2-retro-miRNA-treated and rAAV2-retro-Scramble-treated *SOD1*^G93A^ mice. Differences were assessed using the log-rank (Mantel-Cox) test. (B) Kaplan-Meier survival curves of rAAV2-retro-miRNA-treated and rAAV2-retro-Scramble-treated *SOD1*^G93A^ mice. Differences were assessed using the log-rank (Mantel-Cox) test. (C) Quantification of median survival in rAAV2-retro-miRNA-treated and rAAV2-retro-Scramble-treated *SOD1*^G93A^ mice. Data represent means ± SEM. Student’s t-test. (D) Early disease duration in miRNA-treated and scramble-treated *SOD1*^G93A^ mice. Data represent means ± SEM, Student’s test. (E) Disease duration in rAAV2-retro-miRNA-treated and rAAV2-retro-Scramble-treated *SOD1*^G93A^ mice. Data represent means ± SEM, Student’s t-test. (F) Total disease duration in miRNA-treated and scramble-treated *SOD1*^G93A^ mice. Data represent means ± SEM, Student’s t-test; n = 12 mice per group. \**P* < 0.05, \*\*\**P* < 0.001, \*\*\*\**P* < 0.0001.

Further analysis of disease phases revealed that the early disease period (from peak weight to 10% weight loss) was significantly prolonged in miRNA-treated mice (67.08 days vs. 42.00 days in controls; *P* = 0.0023; Figure 2D). Similarly, the late disease phase, defined as the interval between 10% weight loss and endpoint, was significantly lengthened (25.08 days vs. 16.08 days; *P* = 0.0215; Figure 2E). Consequently, the total disease duration was substantially increased in the rAAV2-retro-miRNA group (92.17 days) relative to the scramble control group (58.08 days; *P* = 0.0002; Figure 2F). These results demonstrate that a single neonatal IM injection of rAAV2-retro-miRNA markedly delays disease progression and improves survival in this ALS model.

### Intramuscular Delivery of rAAV2-Retro-miRNA Preserves Motor Function in *SOD1*^G93A^ Mice

To assess neuromuscular integrity comprehensively, a longitudinal behavioral analysis was conducted beginning at postnatal day 60. The experimental battery included rotarod, grip strength, open-field, and limb clasping assays, all conducted by investigators blinded to the treatment groups. Administration of rAAV2-retro-miRNA markedly preserved motor function. By 18 weeks, open-field locomotor activity declined by only 33.9% in miRNA-treated mice, compared to a 75.74% reduction observed in scramble controls (Figure 3A). Notably, by day 135, all scramble-treated mice exhibited dystonic limb clasping, whereas one-third of miRNA-treated animals remained completely free of this neurological sign (Figure 3B). Furthermore, miRNA-treated mice demonstrated significantly enhanced motor coordination on the rotarod and maintained higher forelimb and hindlimb grip strength compared to scramble controls (Figures 3C and 3D). Collectively, these findings demonstrate that rAAV2-retro-miRNA treatment sustains motor function and muscle strength over prolonged disease progression.

**Figure 3.**
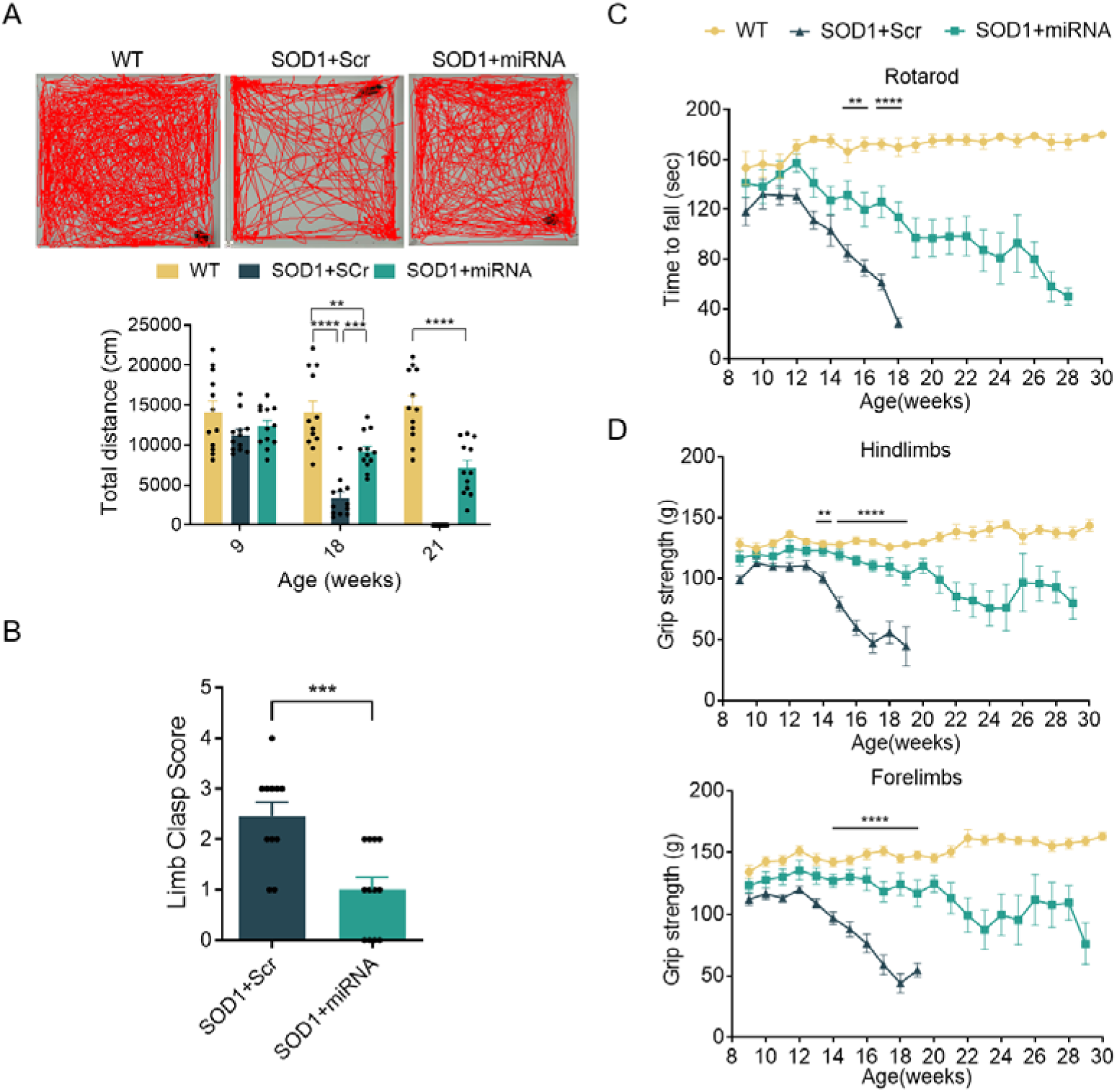
Intramuscular delivery of rAAV2-retro-miRNA leads to long-term preservation of motor function in neonatal *SOD1*^G93A^mice. (A) Open-field motor activity, measured as total distance traveled per hour, in WT, rAAV2-retro-Scramble-treated, and rAAV2-retro-miRNA-treated ALS mice. Scale bar, 10 cm. One-way ANOVA with Tukey’s post hoc test. (B) Limb clasping score in WT, rAAV2-retro-Scramble-treated, and rAAV2-retro-miRNA-treated ALS mice. Student’s t-test. \*\**P* < 0.01, \*\*\**P* < 0.001, \*\*\*\**P* < 0.0001. (C) Rotarod performance of WT, rAAV2-retro-Scramble-treated, and rAAV2-retro-miRNA-treated ALS mice. One-way ANOVA with Tukey’s post hoc test. (D) Forelimb and hindlimb grip strength in WT, rAAV2-retro-Scramble-treated, and rAAV2-retro-miRNA-treated ALS mice. One-way ANOVA with Tukey’s post hoc test. Data represent mean ± SEM; n = 12 mice per group. \*\**P* < 0.01, \*\*\*\**P* < 0.0001 for the comparison between rAAV2-retro-Scramble-treated and rAAV2-retro-miRNA-treated groups.

### rAAV2-Retro-miRNA Gene Delivery Mitigates Neuromuscular Junction Denervation and Muscle Atrophy in *SOD1*^G93A^ Mice

To evaluate the protective effects of rAAV2-retro-miRNA on the integrity of neuromuscular junction (NMJ) and muscle morphology, NMJs and myofibers were systematically analyzed. Consistent with the progressive pathology of ALS, only 10.12% of NMJs remained fully innervated in scramble-treated *SOD1*^G93A^ mice by day 130, with the majority exhibiting partial or complete denervation (Figures 4A and 4B). In contrast, neonatal administration of rAAV2-retro-miRNA significantly preserved NMJ innervation and mitigated denervation.

**Figure 4.**
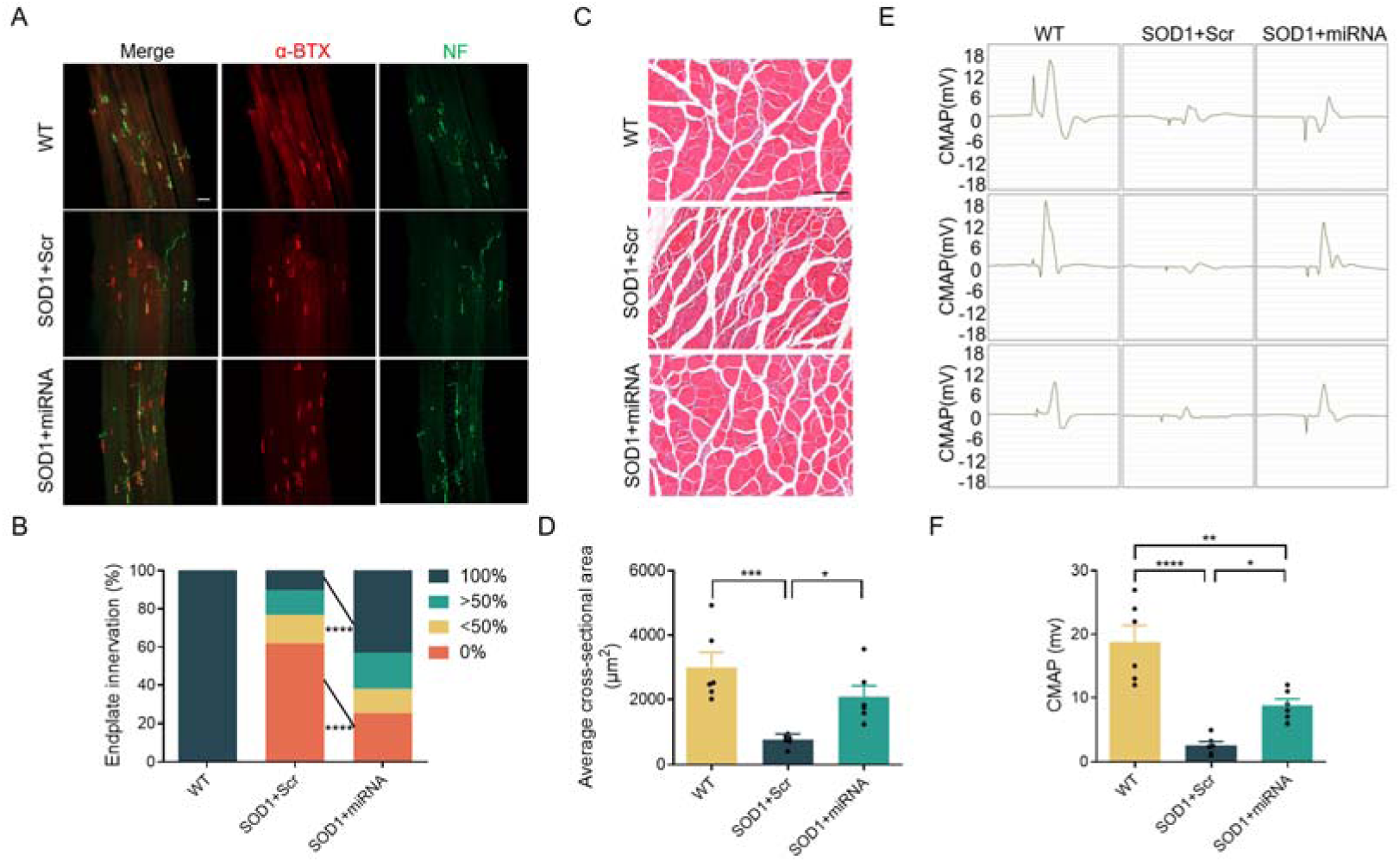
Intramuscular delivery of rAAV2-retro-miRNA preserves neuromuscular junctions, muscle fiber integrity, and motor function in neonatal *SOD1*^G93A^ mice. (A) Representative immunofluorescence images of gastrocnemius muscles from WT, rAAV2-retro-Scramble-treated, or rAAV2-retro-miRNA-treated ALS mice. Sections were stained with antibodies against neurofilament (NF, presynaptic marker) and α-bungarotoxin (α-BTX, postsynaptic acetylcholine receptor marker). Scale bar, 50 µm. (B) Quantification of neuromuscular junction (NMJ) innervation status, categorized as fully innervated (100%), partially innervated, or fully denervated (0%). Student’s t-test. (C) Representative H&E-stained sections of gastrocnemius muscle. Scale bar: 150 μm. (D) Mean cross-sectional area (CSA) of muscle fibers. One-way ANOVA with Tukey’s post hoc test. (E) Representative compound muscle action potential (CMAP) recordings from wild-type non-transgenic (left), rAAV2-retro–Scramble-treated (middle), and rAAV2-retro-miRNA-treated *SOD1^G93A^* mice (right). (F) Quantitative analysis of CMAP amplitudes. One-way ANOVA with Tukey’s post hoc test. Data represent mean ± SEM, n=6 mice per group. \**P* < 0.05, \*\**P* < 0.01, \*\*\**P* < 0.001, \*\*\*\**P* < 0.0001.

Given the critical role of NMJ integrity in maintaining muscle mass, myofiber cross-sectional area(CSA) was quantified using laminin immunostaining. Scramble-treated *SOD1*^G93A^ mice displayed pronounced muscle atrophy, with a mean CSA (765.7 μm²) significantly smaller than that observed in wild-type mice (3009 μm²; *P* = 0.0008; Figures 4C and 4D). Treatment with rAAV2-retro-miRNA markedly attenuated atrophy, increasing mean CSA to 2090 μm² (*P* = 0.0038 *vs.* scramble; Figures 4C and 4D), and preventing whole-muscle mass loss (Figure S3).

Functional assessment of NMJ transmission at 130 days revealed significantly higher CMAP amplitudes in rAAV2-retro-miRNA-treated mice compared to scramble controls (Figures 4E and 4F), indicating improved neuromuscular transmission and delayed functional denervation. Collectively, these results demonstrate that rAAV2-retro-miRNA therapy effectively preserves NMJ innervation, alleviates muscle atrophy, and maintains electrophysiological function in *SOD1*^G93A^ mice.

### Intramuscular Delivery of rAAV2-Retro-miRNA Preserves Spinal α-motoneurons and Suppresses Mutant SOD1 Expression in *SOD1*^G93A^ Mice

Degeneration of spinal motor neurons and skeletal muscle denervation represent key pathological hallmarks of ALS. Immunohistochemical analyses revealed severe loss of α-motoneurons across all spinal cord segments in scramble-treated *SOD1*^G93A^ mice at day 130, compared to wild-type controls. In contrast, treatment with rAAV2-retro-miRNA significantly preserved α-motoneuron survival throughout the spinal cord (Figures 5A and 5B).

**Figure 5.**
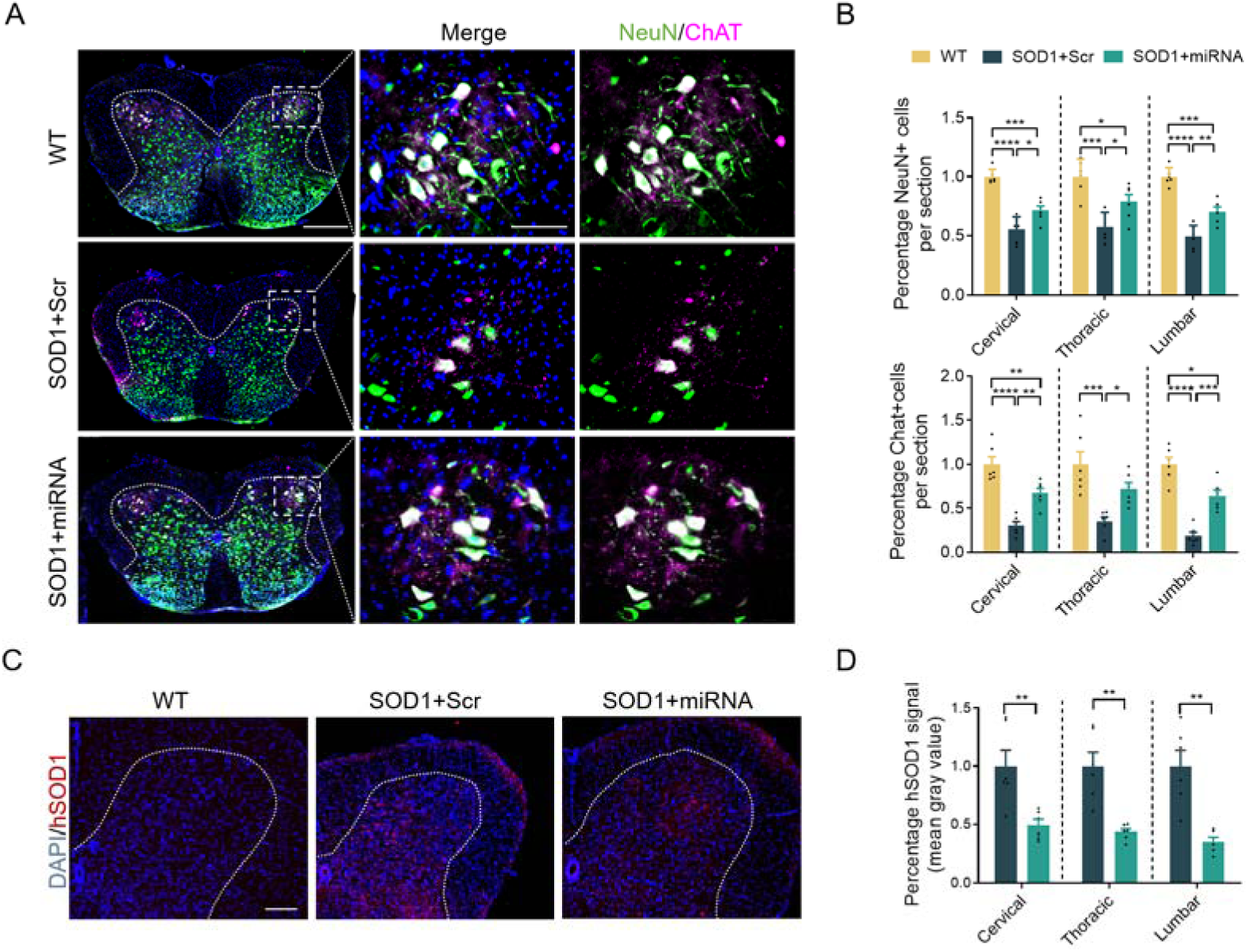
Intramuscular delivery of rAAV2-retro-miRNA preserves α-motor neurons and reduces SOD1 protein accumulation in *SOD1*^G93A^mice. (A) Representative immunofluorescence images of lumbar spinal cord sections from WT, rAAV2-retro-Scramble-treated, and rAAV2-retro-miRNA-treated ALS mice at day 130. Staining markers: NeuN (green), choline acetyltransferase (ChAT, carmine). Nuclei are counterstained with DAPI (blue). Scale bar, 400 µm(left) or 100 µm(right). (B) Quantification of α-motor neurons and NeuN^+^ neurons in cervical, thoracic, and lumbar spinal cord regions. One-way ANOVA with Tukey’s post hoc test. (C) Representative images of hSOD1 immunostaining in lumbar spinal cord sections from WT, rAAV2-retro-Scramble-treated, and rAAV2-retro-miRNA-treated ALS mice on day 130. Nuclei are labeled with DAPI (blue); hSOD1 is shown in red. Scale bar, 200 µm. (D) Quantitative analysis of h*SOD1* expression levels in cervical, thoracic, and lumbar spinal cord regions. Student’s t-test. Data represent mean ± SEM n=6 mice per group. \**P* < 0.05, \*\**P*< 0.01, \*\*\**P* < 0.001, \*\*\*\**P* < 0.0001.

Quantitative analysis demonstrated that scramble-treated *SOD1*^G93A^ mice exhibited a decline in NeuN neuronal count to roughly 50% of wild-type levels (cervical: 55.44%; thoracic: 57.44 %; lumbar: 48.94%), whereas rAAV2-retro-miRNA treatment maintained NeuN neurons above 70% of normal values (cervical: 76.16%; thoracic: 74.49%; lumbar: 70.32%). Consistent with these neuroprotective effects, pronounced hSOD1 immunoreactivity was observed in the spinal cord of scramble-treated *SOD1*^G93A^ mice, while rAAV2-retro-miRNA administration markedly suppressed hSOD1 expression (Figures 5C and 5D).

Collectively, these findings demonstrate that neonatal IM delivery of rAAV2-retro-miRNA effectively downregulates mutant SOD1 levels and provides broad neuroprotection across the spinal cord.

### Intramuscular Delivery of rAAV2-Retro-miRNA Attenuates Neuroinflammation in the Spinal Cord of *SOD1*^G93A^ Mice

Neuroinflammation, characterized by the activation of microglia and astrocytes, is a prominent pathological hallmark of ALS. To determine whether rAAV2-retro-miRNA treatment modulates neuroinflammatory response, immunofluorescence analyses were performed on spinal cord sections. Treatment with rAAV2-retro-miRNA markedly reduced the fluorescence intensity of both Iba1 (microglia) and GFAP (astrocytes) relative to scramble-injected *SOD1*^G93A^ controls (Figure 6), indicating suppression of glial activation. These results demonstrate that targeted silencing of mutant *SOD1* via rAAV2-retro-miRNA not only preserves motor neurons but also attenuates neuroinflammation in this ALS model.

**Figure 6.**
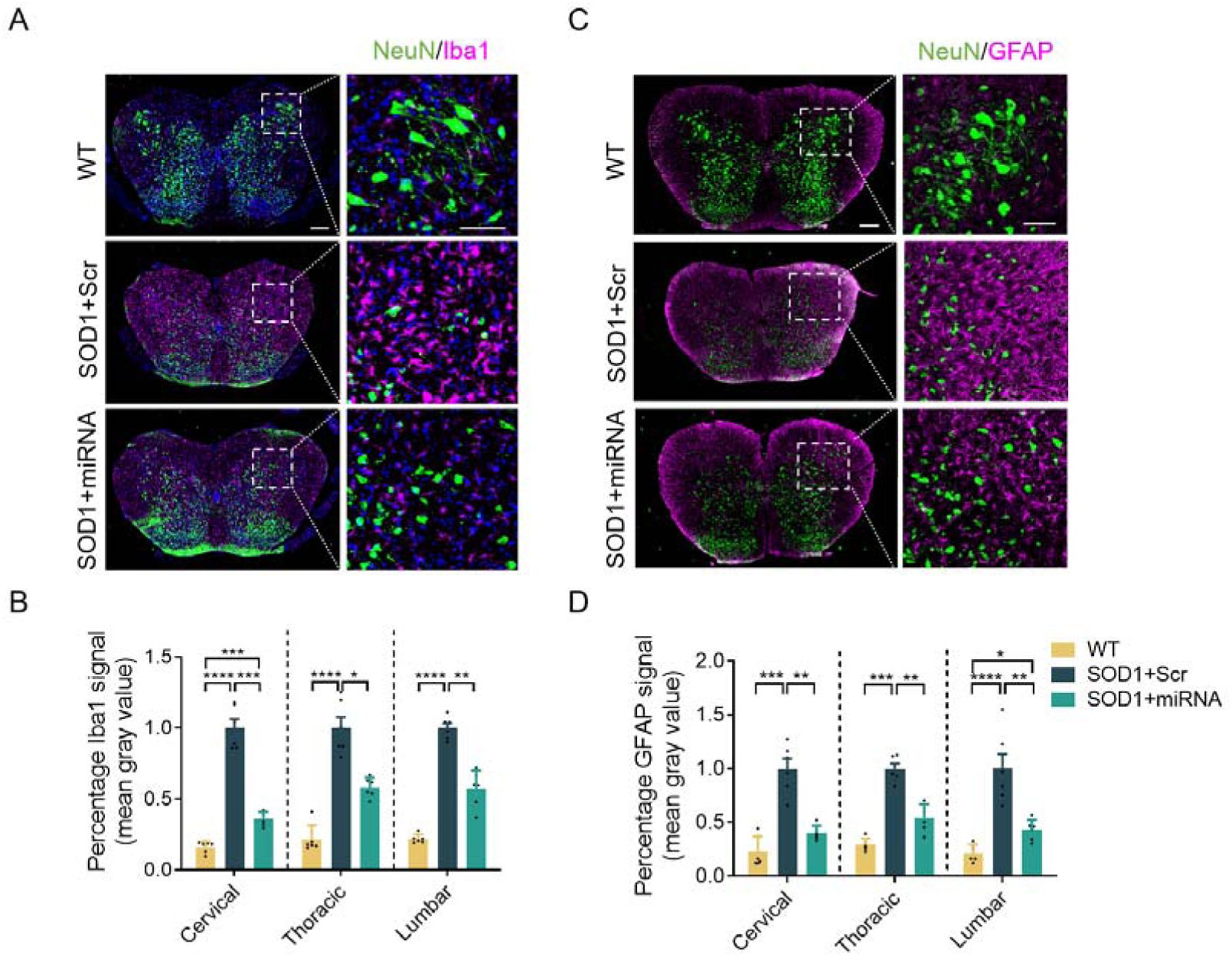
Intramuscular delivery of rAAV2-retro-miRNA attenuates neuroinflammation in *SOD1*^G93A^ mice. (A)Representative immunofluorescence images of Iba1^+^ microglia in lumbar spinal cord sections from WT, rAAV2-retro-Scramble-treated, and rAAV2-retro-miRNA-treated ALS mice. Scale bars, 200 µm (left) or 100 µm (right). (B) Quantification of Iba1 fluorescence intensity in the gray matter of wild-type (WT) and treated mice. (C) Representative immunofluorescence images of GFAP^+^ astrocytes in lumbar spinal cord sections from WT, rAAV2-retro-Scramble-treated, and rAAV2-retro-miRNA-treated ALS mice. Scale bars, 200 µm (left) or 100 µm (right). (D) Quantification of GFAP fluorescence intensity in the gray matter. Data represent mean ± SEM, n=6 mice per group. One-way ANOVA with Tukey’s post hoc test. \**P* < 0.05, \*\**P* < 0.01, \*\*\**P* < 0.001, \*\*\*\**P* < 0.0001.

## Discussion

This study establishes a gene delivery platform that integrates the retrograde transport capability of rAAV2-retro with the safety and precision of an artificial miRNA to achieve targeted silencing of the human SOD1 (h*SOD1*) gene. A single IM administration of rAAV2-retro-miRNA in neonatal *SOD1*^G93A^ mice led to widespread transgene expression throughout the spinal cord, significant suppression of mutant SOD1 protein, and a broad spectrum of therapeutic benefits, including delayed disease onset, extended lifespan, improved motor function, and preservation of NMJs and spinal motor neurons. These findings provide a proof-of-concept validation for the rAAV2-retro-miRNA platform as a versatile tool for spinal motor neuron-targeted gene silencing.

The application of rAAV2-retro, a vector exhibiting superior retrograde transduction efficiency in neonatal animals, addresses the limited spinal cord transduction observed with conventional AAV serotypes in adults. rAAV2-retro was originally developed through directed evolution to enhance retrograde transport from axon terminals to neuronal cell bodies. ^32^Unlike AAV9, which often shows prominent transduction of astrocytes when delivered peripherally, ^33^rAAV2-retro exhibits a preferential tropism for motor neurons via retrograde axonal transport. Neonatal IM administration of rAAV2-retro-CAG-miRNA achieved robust and widespread gene silencing across spinal motor neurons, surpassing the restricted expression patterns reported in prior studies.

In addition to the choice of vector, the route of administration is a critical determinant of transduction efficiency and safety. Current gene delivery strategies for ALS include intrathecal (IT), intracerebroventricular (ICV), and intraparenchymal injections, all of which require invasive procedures and pose safety concerns.^38,39^ In contrast, intramuscular (IM) injection provides a peripheral route for vector administration. Studies have shown that IM injection of rAAV vectors enables retrograde axonal transport to spinal motor neurons, resulting in CNS transduction with limited systemic exposure. For example, in *SOD1*^G93A^ ALS mouse models, IM delivery of AAV6-HGF improved muscle histology and delayed disease progression,^28^ and IM delivery of AAV9-IGF2 preserved motor neurons and promoted axonal regeneration.^26^ These observations support the feasibility of using IM injection as a peripheral route for motor neuron-targeted gene delivery.

Mechanistically, our strategy differs from prior IM studies. Earlier approaches primarily delivered neurotrophic factors (e.g., IGF-1, IGF-2, HGF) via IM injection to ameliorate pathology. While beneficial, these approaches offer modest efficacy due to their single-mechanism, symptomatic focus.^24–26, 28^ In contrast, the present study employs rAAV2-retro-mediated *SOD1* gene silencing during the neonatal period, directly addressing the genetic etiology of ALS, thereby preventing the accumulation of toxic protein and mitigating downstream pathogenic mechanisms such as mitochondrial dysfunction and neuroinflammation. This “upstream interception” strategy targets the core pathology of ALS, offering a mechanistic advantage over neuroprotective interventions that primarily enhance neuronal survival without addressing the underlying cause. By directly intervening at the primary pathogenic source, this causal approach yields multidimensional therapeutic benefits, including improved motor function, enhanced motor neuron survival, reduced apoptosis and neuroinflammation, preserved NMJs, and mitigation of muscle atrophy.

The neonatal period offers several biological features that make it a well-defined system for platform validation. During this developmental window, prior to the onset of motor neuron synaptic stripping^34^ and while the complement system remains immature,^35^ the favorable permeability properties of the BBB facilitates efficient CNS transduction following IM injection of rAAV2-retro. In addition, the immaturity of the neonatal immune system diminishes adaptive immune responses against both the viral capsid and the transgene product, thereby promoting sustained and stable transgene expression. Leveraging these features, a single IM dose of rAAV2-retro-miRNA achieved sustained transgene expression and prolonged efficacy throughout postnatal development, consistent with the genomic stability of rAAV2-retro and the favorable immunological context of the neonatal period. In this pre-symptomatic *SOD1*^G93A^ model, neonatal administration achieved pan-spinal *SOD1* silencing, effectively disrupting the pathogenic feed-forward loop between neuroinflammation and NMJ degeneration and markedly delaying disease progression. This finding aligns with a previous study showing that combined ICV/IV administration at postnatal day 1 also extended survival in the same model,^34^ suggesting that early postnatal intervention can be effective in this preclinical setting. Together, these results highlight the utility of the neonatal period as a system for evaluating platform efficacy and have broader implications for motor neuron-targeted gene delivery.

Consistent with this notion, the neonatal intramuscular route has been employed in previous studies for gene delivery to motor neurons. A landmark study using an EIAV lentiviral vector achieved robust *SOD1* silencing and extended survival in the same ALS mouse model following neonatal intramuscular injection.^35^ More recently, intramuscular delivery of scAAV9 in neonatal SMA mice mediated widespread spinal cord transduction and markedly extended lifespan,^36^ demonstrating the feasibility of this route for motor neuron-targeted gene delivery. In the context of ALS, however, AAV-mediated miRNA delivery via the neonatal intramuscular route has not been previously reported. Our study thus provides a platform validation for the rAAV2-retro-miRNA platform in this setting.

It is noteworthy that the previous EIAV-based study^35^ validated the feasibility of peripheral RNAi delivery for *SOD1*-ALS, yet the present study extends this paradigm in two important aspects. First, we employed an AAV2-retro vector, which offers sustained transgene expression without genomic integration, addressing a key concern associated with lentiviral vectors. Second, we utilized an artificial miRNA instead of shRNA, which has been shown to mitigate the risk of RNAi pathway oversaturation and associated toxicity. ^22, 23^ Thus, while both studies establish the therapeutic potential of peripheral gene silencing, the AAV-miRNA platform described here offers a distinct safety advantage for motor neuron-targeted gene delivery.

Mutations in the superoxide dismutase 1 (*SOD1*) gene account for approximately 20% of familial ALS cases, with over 150 pathogenic variants identified to date.^37^ Substantial evidence indicates that mutant SOD1 acquires toxic gain-of-function properties, leading to protein misfolding and aggregation, a histopathological hallmark of *SOD1*-linked ALS.^38, 39^ Notably, misfolded *SOD1* inclusions have also been detected in a subset of sporadic ALS cases.^40, 41^ Although the precise mechanistic cascade remains incompletely elucidated, converging evidence supports that *SOD1*-mediated toxicity perturbs critical cellular functions, collectively driving motor neuron degeneration.^42, 43^ Consequently, suppression of mutant *SOD1* expression represents a rational therapeutic strategy. Preclinical studies have consistently demonstrated that silencing mutant *SOD1* prolongs survival, delays symptom onset, and preserves motor function in ALS rodent models, thereby establishing *SOD1* as a well-validated target for gene therapy development.^44^ Unlike neurotrophic support strategies (e.g., IGF1, BDNF) that modulate downstream consequences of neurodegeneration, direct *SOD1* silencing intervenes at the primary pathogenic source.^17^ As the first genetically defined form of ALS, *SOD1* has been extensively investigated in AAV-RNAi-based interventions, with progressive refinement of therapeutic efficacy.^45^ The present study builds on this foundation by employing a rAAV2-retro IM platform to achieve efficient and widespread motor neuron transduction and *SOD1* silencing in neonatal animals. This platform may also be adaptable for targeting other ALS-linked genes (e.g., *C9orf72, FUS, TARDBP*)^46^ or motor neuron disease-associated mutations.

Beyond the present platform validation, this platform has several potential directions for further development. Future directions include designing dual-component vectors that co-deliver *SOD1*-targeting miRNAs with neuroprotective factors such as IGF1 in subsequent studies, as well as developing combinatorial regimens integrating neonatal gene silencing with adult-stage therapies. The pan-spinal motor neuron transduction achieved by rAAV2-retro via IM injection unlocks potential for spinal cord research. Beyond ALS, this platform facilitates precise circuit dissection using Cre/loxP or CRISPR systems, facilitating targeted gene manipulation in models of hereditary spastic paraplegia or spinal muscular atrophy. Integration with optogenetic tools (e.g., ChR2) or chemogenetic systems (e.g., DREADDs) permits real-time functional mapping of motor ensembles involved in locomotion and pain processing, while calcium indicators (e.g., GCaMP) link histological alterations to functional outcomes during disease progression. The approach also supports multi-target interventions that combine neuroprotection, gene silencing, and regeneration. Continued optimization of vector tropism in relevant models will further expand the utility of this platform for spinal and motor neuron research.

## Methods

### Animals

C57BL/6J (stock no.000664) and high-copy transgenic *SOD1* ^G93A^ mice [B6. Cg-Tg(*SOD1*^G93A^)/1Gur/J] (stock no.004435) were obtained from The Jackson Laboratory (Bar Harbor, ME, USA). Transgenic littermates of both sexes expressing elevated levels of the mutant *SOD1*^G93A^ transgene were randomly and equally assigned to experimental groups, while non-transgenic littermates served as wild-type controls. All animals were maintained under specific pathogen-free (SPF) conditions in the institutional animal facility with controlled temperature, humidity, light, and noise levels and kept on a 12-hour light/dark cycle with *ad libitum* access to food and water. All experimental procedures were approved by the Animal Experimental Ethics Committee of Shanghai Medical School, Fudan University, China (protocol code 2024-DLAS-001), and were performed in accordance with the institutional guidelines.

### Production, Purification, and Titration of Recombinant AAV Vectors

A single-stranded recombinant AAV2-retro vector was engineered to express enhanced green fluorescent protein (eGFP) under the control of a CAG promoter. The construct also incorporated a synthetic miRNA sequence designed to target h*SOD1* mRNA (exon 2), followed by the woodchuck hepatitis virus post-transcriptional regulatory element (WPRE) and an SV40 polyadenylation (poly(A)) signal. The artificial miRNA, optimized to specifically silence h*SOD1*without cross-reactivity to mouse *Sod1*, was embedded within the human *miR-30a* scaffold, as previously described.^31^ The complete oligonucleotide sequence of the artificial miRNA is provided below, with the guide strand underlined: miR-30a construct:5’-TGTTTGAATGAGGCTTCAGTACTTTACAGAATCGTTGCCTGCACA TCTTGGAAACACTTGCTGGGATTACTTCTTCAGGTTAACCCAACAGAAGGCAAAGAAG GTATATTGCTGTTGACAGTGAGCG<u>TCATGAACATGGAATCCATG</u>TAGTGAAGCCACAGA TGTAAAGGTGGATGAAGAAAGTATGCCTACTGCCTCGGACTTCAAGGGGCTACTTTAG GAGCAATTATCTTGTTTACTAAAACTGAATACCTTGCTATCTCTTTGATACATTTTTACAA AGCTGAATTAAAATGGTATAAATTAAATCACTTTA-3’.

Recombinant vectors were packaged into AAV2 capsids using a triple-plasmid transfection in HEK 293 cells. The transfection mixture comprised: (1) the AAV2 inverted terminal repeat (ITR)-based transgene plasmid, (2) a helper plasmid expressing AAV2 *Rep* and *Cap* genes, and (3) the adenoviral helper plasmid pHELP. Subsequently, viral particles were purified, and titers were determined using quantitative PCR (qPCR).

### Intramuscular Injection

As described previously, cryoanesthesia was induced in postnatal day 4 (P4) mice by briefly placing them on pre-chilled aluminum plates.^29^ For IM administration, small incisions were made using fine surgical scissors to expose target muscles (the left flexor carpi ulnaris and the right gastrocnemius). Recombinant AAV vectors were diluted in phosphate-buffered saline (PBS) to a concentration of 1 × 10¹³ genomic copies (GC)/mL. A total of 2 µL, containing 2 × 10^10^ GC of each vector, was injected into each muscle using a glass micropipette connected to a high-precision syringe pump (Quintessential Stereo taxic Injector, Model 53311). Injections were administered at a constant rate of 1.0 μL/min. To minimize reflux and ensure complete delivery, the pipette was held in place for 30 seconds following each injection before being slowly withdrawn. After the procedure, pups were allowed to recover on a warming pad before being returned to their home cages.

### Western Blot

Mice were deeply anesthetized, decapitated, and tissues were rapidly dissected, snap-frozen in liquid nitrogen, and stored at -80°C. For protein extraction, frozen tissues or cultured cells were homogenized in RIPA buffer (Beyotime, P0013B) supplemented with a protease inhibitor cocktail (Beyotime, ST506). Lysates were mixed with loading buffer (Tsingke, TSJ010) and boiled at 100°C for 10 minutes. Equal amounts of proteins were separated by SDS-PAGE and transferred onto polyvinylidene fluoride (PVDF) membranes. Membranes were blocked with 5% non-fat dry milk in Tris-buffered saline containing 0.05% Tween-20 (TBST; 20 mM Tris-HCl, 150 mM NaCl, pH 7.5) for 1 hour at room temperature, followed by overnight incubation at 4°C with primary antibodies. After washing, membranes were incubated with horseradish peroxidase (HRP)-conjugated secondary antibodies. The following antibodies were used: mouse anti-hSOD1 (1:200, Santa Cruz Biotechnology, sc-17767), mouse anti-Actin (1:5000, Proteintech, 66009-1-IG), mouse anti-eGFP (1:1000, Santa Cruz Biotechnology, sc-5384), and HRP-conjugated goat anti-mouse IgG (1:5000, YEASEN, 33201ES60). Protein bands were visualized using enhanced chemiluminescence, and band intensities were quantified using Image Lab (Bio-Rad) or ImageJ software, normalized to β-Actin levels in each lane.

### RNA Extraction, Reverse Transcription, and Quantitative Real-Time PCR

Total RNA was isolated from cultured N2A cells using TRIzol™ Reagent (Thermo Fisher Scientific, 15596018CN) as per the manufacturer’s protocol. RNA concentration and purity were determined spectrophotometrically by measuring absorbance at 260 and 280 nm, and integrity was confirmed by the 260/280 nm ratio. Complementary DNA (cDNA) was synthesized from 1 μg of total RNA using random primers and the Hifair® III First Strand cDNA Synthesis Kit (Yeasen Biotechnology, 11141ES60).

qPCR was performed using the 2× SYBR Green qPCR Master Mix (Selleck Chemicals, B21203) on a Bio-Rad iCycler iQ6 Detection System. Each reaction contained cDNA corresponding to 100 ng of total RNA. Amplification parameters followed the manufacturer’s instructions. The human h*SOD1* gene was amplified with the following primers:

Forward: 5′-GGGAAGCTGTTGTCCCAAG-3′

Reverse: 5′-CAAGGGGAGGTTAAAAGAGAGC-3′

Mouse GAPDH (NM_001289726.2) served as endogenous control and was amplified with the following primers:

Forward: 5′-CACCATCTTCCAGGAGCGAG-3′

Reverse: 5′-CCTTCTCCATGGTGGTGAAGAC-3′

Relative mRNA expression levels were calculated using the 2^−ΔΔCt^ method, and data acquisition and analysis were performed with Bio-Rad iQ6 Optical System Software.

### Behavioral Analysis

Behavioral assessments were conducted to evaluate disease progression and motor performance in wild-type (WT), rAAV2-retro-Scramble-treated, and rAAV2-retro-miRNA-treated *SOD1* ^G93A^ mice. All tests were performed during the light phase under consistent environmental conditions, and experimenters were blinded to group identity throughout data collection.

#### Body Weight Measurement

Body weight was recorded weekly beginning at postnatal day 60 and continued until end-stage disease to monitor disease progression.

#### Rotarod Test

Motor coordination and balance were assessed using an accelerating rotarod apparatus (KEW BASIS, KW-6D). Mice underwent three consecutive trials per session, with the rotation speed gradually increasing from 5 rpm to 40 rpm over a minute. The latency to fall was recorded for each trial.

#### Limb Clasping Assay

Hindlimb clasping behavior was evaluated at 135 days of age by suspending mice by the tail for 10 seconds with the heads oriented downward. A clasping score was assigned if the behavior persisted for more than 2 seconds using the following criteria: 1 point for unilateral hindlimb clasping, 2 points for bilateral hindlimb clasping, and 3 points if clasping involved both hindlimbs, forelimbs, or torso curling.

#### Grip-Strength Test

Forelimb and hindlimb grip strength were quantified using a grip-strength meter (KEW BASIS, KW-ZL). Mice were allowed to grasp a horizontal metal bar with either forelimbs or hindlimbs, and were gently pulled backward in a consistent motion until they released the bar. The peak tension (in grams) was recorded over five consecutive trials, and the mean value was calculated for each mouse.

#### Open-Field Test

Spontaneous locomotor activity was evaluated in a clean, opaque acrylic open-field arena (50 cm × 50 cm) for one hour. Animal movement was recorded and analyzed using EthoVision Video Tracking Software (Noldus). The total distance traveled and movement patterns were used to analyze locomotor activity and exploratory behavior.

### Assessment of Disease Progression and Survival

The humane experimental endpoint was defined as the time point at which an animal was unable to right itself within 30 seconds of being placed on its back. Disease onset and progression were evaluated retrospectively based on recorded physiological and behavioral data. Disease onset was defined as the age at which animals sustained a 10% loss from their peak body weight. The disease course was further subdivided into two phases: the early phase, extending from peak body weight to a 10% reduction in peak weight, and the late phase, spanning from 10% weight loss threshold to the endpoint.

### Compound Muscle Action Potential (CMAP) Measurement

At 130 days of age, mice were anesthetized with isoflurane, and hair from the hindlimb and lower back regions was gently removed using depilatory cream before electrophysiological recording. The stimulating electrode was positioned at the lumbar spinal roots, while the active recording electrode was placed over the belly of the gastrocnemius muscle. The reference electrode was placed near the Achilles tendon or digit of the ipsilateral hindlimb (iWork, IX-RA-834ECG).

### Tissue Preparation

Following euthanasia via isoflurane overdose, mice were transcardially perfused with phosphate-buffered saline (PBS) to remove circulating blood, followed by fixation with 4% paraformaldehyde (PFA) in PBS. The perfused tissues were post-fixed in 4% PFA overnight at 4°C. Spinal cords were cryoprotected in 30% sucrose for 24 hours and sectioned at a thickness of 30 μm using a freezing microtome (Leica, CM3050 S). Gastrocnemius muscle and liver tissues were dehydrated, embedded in paraffin, and sectioned into 3 μm slices using a rotary microtome (Leica, 149BIO000C1).

### Immunohistochemistry and Histological Analysis

For immunohistochemical staining of spinal cord sections, fixed sections were rinsed three times with PBS and subjected to antigen retrieval using 1× citrate buffer (Sangon Biotech, A100101-0500). Sections were then permeabilized and blocked for one hour at room temperature with 5% goat serum (Sangon Biotech, E510009-0100) and 0.3% Triton X-100 (Beyotime, P0096) in PBS. Following blocking, samples were incubated overnight at 4°C with primary antibodies diluted in PBS containing 3% goat serum (Sangon Biotech, E510009-0100) and 0.1% Triton X-100 (Beyotime, P0096). The following primary antibodies were used: rabbit anti-ChAT (1:1000, Abcam, ab178850), chicken anti-NeuN (1:1000, Millipore, ABN91), mouse anti-eGFP (1:1000, Santa Cruz Biotechnology, sc-5384), rabbit anti-Iba1 (1:1000, Abcam, ab178846), rabbit anti-GFAP (1:1000, Abcam, ab7260), rabbit anti-hSOD1 (1:500, Cell Signaling Technology, 2770).

After washing three times with PBS, sections were incubated for 2 hours at room temperature in the dark with appropriate secondary antibodies: donkey anti-mouse Alexa Fluor 488 (1:500, Thermo Fisher, A-20102), goat anti-chicken Alexa Fluor 647 (1:1000, Abcam, ab150171), and donkey anti-rabbit Alexa Fluor 555 (1:500, Thermo Fisher, A-31572). Nuclei were counterstained with DAPI (1:1000, Sigma-Aldrich, 10236276001) for 10 minutes, followed by three PBS washes. Sections were mounted using antifade mounting medium (SouthernBiotech, 0100-01) prior to imaging.

For immunohistochemical staining of liver sections, paraffin-embedded liver tissues were deparaffinized in xylene (Sinopharm Chemical Reagent, 10023428) and rehydrated through a graded ethanol series (Sinopharm Chemical Reagent, 100092680). Antigen retrieval was performed using microwave heating in Tris-EDTA buffer (Solarbio, C1038). Sections were then blocked with 5% normal goat serum for 1 hour at room temperature and incubated overnight at 4°C with the following primary antibody: mouse anti-eGFP (1:1000, Santa Cruz Biotechnology, sc-5384). Following three washes with TBS-T, slides were incubated for 1–2 hours at room temperature in the dark with the corresponding secondary antibody: donkey anti-mouse Alexa Fluor 488 (1:500, Thermo Fisher Scientific, A-20102). Nuclei were counterstained with DAPI (1:1000, Sigma-Aldrich, 10236276001) for 10 minutes, washed with TBS-T, and mounted with antifade mounting medium (SouthernBiotech, 0100-01) for subsequent imaging.

For NMJ immunostaining, gastrocnemius muscles were dissected and teased into small bundles containing 2-3 fibers, as previously described^47^. Muscle bundles were incubated with chicken anti-neurofilament (1:2500, Abcam, ab4680) to label presynaptic axons and fluorophore-conjugated α-bungarotoxin (α-BTX, Biotium, 00018) to identify postsynaptic acetylcholine receptors.

For H&E Staining, paraffin-embedded gastrocnemius muscle sections were deparaffinized in xylene (Sinopharm Chemical Reagent, 10023428) and rehydrated through a graded ethanol series (Sinopharm Chemical Reagent, 100092680). Nuclei were stained with Harris hematoxylin (Beyotime, C0107) for 5 minutes, rinsed under tap water, and differentiated in 1% acid alcohol (Beyotime, C0163M). Sections were blued in Scott’s tap water substitute and counterstained with eosin (Beyotime, C0109) for 1 minute. After dehydration and xylene clearing, slides were mounted with a resinous medium (Sangon, E675007) for histopathological evaluation.

### Fluorescence Microscopy and Imaging Analysis

Spinal cord and liver sections were initially scanned using a high-resolution digital slide scanner (PANNORAMIC 250 Flash III, 3DHISTECH) equipped with a 20× objective lens. Representative high-magnification images were acquired using a Leica SP8 confocal microscope with a 40× oil-immersion objective. Z-stack images were captured to obtain optical sections at high spatial resolution, consisting of 10–15 planes with step intervals of 0.3 μm for liver and 4 μm for spinal cord.

For NMJ imaging, fluorescence acquisition was performed using an Olympus spinSR confocal microscope with a 20× objective lens. Maximum-intensity projection images were generated from z-stacks using Fiji (ImageJ) software for enhanced visualization. Additionally, H&E-stained sections of gastrocnemius muscle were digitized using the PANNORAMIC 250 Flash III slide scanner with a 10× objective.

### Motoneuron Quantification

Motoneuron analysis was conducted using QuPath and Imaris software. TIFF images were first imported into QuPath, where regions of interest (ROIs) were manually delineated. Nuclei within each ROI were identified in the DAPI channel using the built-in “Cell Detection” function to generate initial detection outputs. These detections were further classified using the “Classify” tool, with parameters optimized to accurately distinguish ChAT-positive and NeuN-positive motoneurons. The classifications were cross-validated through manual inspection in Fiji to ensure counting accuracy.

### Quantification of Glial and hSOD1 Signal Intensity

Fluorescence intensity was quantified using ImageJ software. Microscopy images were first imported and converted to 8-bit format to standardize pixel intensity values. For multi-channel datasets, individual channels were separated to isolate specific fluorescent signals. ROIs were delineated based on fluorescence distribution. Intensity thresholds were then carefully adjusted relative to background intensity to enhance measurement accuracy. The “Area” and “Integrated Density” within these ROIs were measured to obtain quantitative values for fluorescence intensity. **Quantification of Neuromuscular Junctions (NMJs)**

Z-stack images of NMJs were acquired and processed using ImageJ as previously described.^47^ Endplates were classified as innervated if they exhibited co-localization of BTX and NF signals. Conversely, endplates showing only BTX staining without an associated NF signal were classified as denervated.

### Cross-sectional area (CSA) of Muscle

CSA of the gastrocnemius muscle was quantified using *Fiji* software. Muscle fibers were first automatically detected using the built-in segmentation tools, followed by manual correction to ensure accurate boundary delineation. The CSA of individual fibers was then calculated using the “Analyze Particles” function. For each mouse, a minimum of 1,000 fibers was analyzed, and the mean CSA was used for subsequent statistical evaluation.

### Statistical Analyses

Statistical analyses were conducted using GraphPad Prism 7 (GraphPad Software, San Diego, CA, USA). Data are presented as means ± SEM for continuous variables. For comparisons involving only two groups, including the molecular and histological analyses of hSOD1 and NMJ, as well as the limb clasping assay and disease phase quantifications (Figures 2C-2F), an unpaired Student’s t-test was used. For multi-group comparisons (e.g., involving WT, SOD1+Scr, and SOD1+miRNA), normality was assessed using the Shapiro-Wilk test and variance homogeneity using Levene’s test. Both assumptions were met, and one-way ANOVA followed by Tukey’s post hoc test was used. Survival analysis was pre-specified as the principal confirmatory endpoint; survival outcomes were visualized by Kaplan–Meier curves and compared using the log-rank (Mantel–Cox) test. All other endpoints were analyzed as exploratory, and P values for these endpoints were not adjusted for multiplicity across the full family of endpoints. For behavioral outcomes (rotarod, grip strength, and open-field test), cross-sectional comparisons were performed at each time point using one-way ANOVA followed by Tukey’s post hoc test. Mixed-effects models were not used because the missing data at late time points were not missing at random, as they resulted from disease progression and animals reaching humane endpoints; this limits the validity of conventional longitudinal inference. Statistical significance was assigned at P < 0.05, with thresholds indicated as *P < 0.05, **P < 0.01, ***P < 0.001, and ****P < 0.0001. The number of replicates per experiment is provided in the corresponding figure legends.

## ACKNOWLEDGMENTS

This work was sponsored by the Natural Science Foundation of Shanghai (Grant No. 22ZR1414500 and 21ZR1407700 to T.X.), and supported by the National Key Research and Development Program of China (Grant No. 2024YFA1108000), the National Natural Science Foundation of China (Grant No. 82441053), the Shanghai Key Laboratory of Aging Studies (Grant No. 19DZ2260400), and the Shanghai Municipal Science and Technology Major Project (Grant No. 2019SHZDZX02). We also thank the staff members of the Integrated Laser Microscopy System at the National Facility for Protein Science in Shanghai for their technical support and assistance in data collection and analysis.

## AUTHOR CONTRIBUTIONS

Conceptualization: W.W., T.X.; data curation: X.G., H.W., Q.W.,J.W.,Z.B.; supervision: W.W., T.X.; visualization: X.G., T.X.; writing-original draft: X.G., Y.X., W.W., T.X.

## Supplemental information

**Figure S1:**
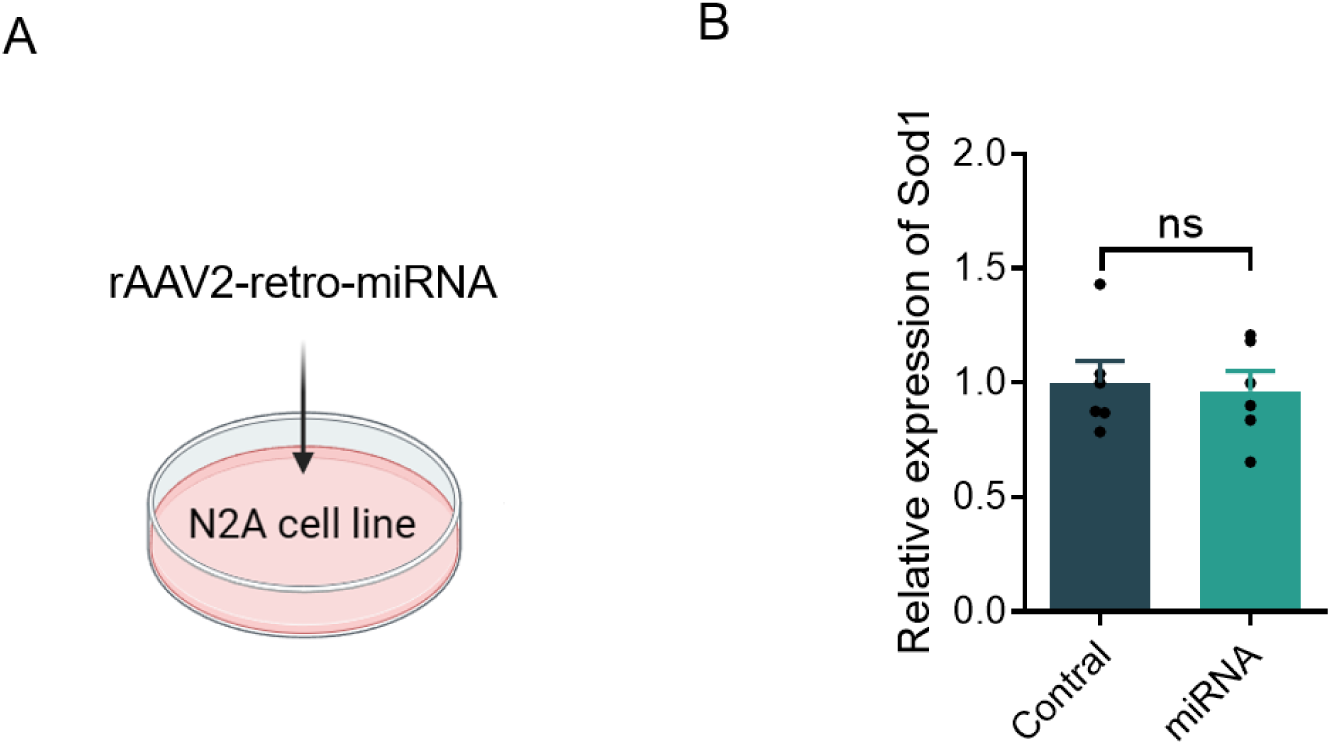
Artificial miRNA does not affect murine SOD1 mRNA expression *in vitro*. (A) N2A cells were transduced with rAAV2-retro–miRNA for 48 hours, followed by total RNA extraction. (B) Quantification of murine *Sod1* (m*Sod1*) mRNA levels via qPCR, normalized to m*GAPDH*. Data are presented as individual biological replicates (dots). ns, not significant. Data represent mean ± SEM, n=6 per group. Student’s t-test.

**Figure S2 :**
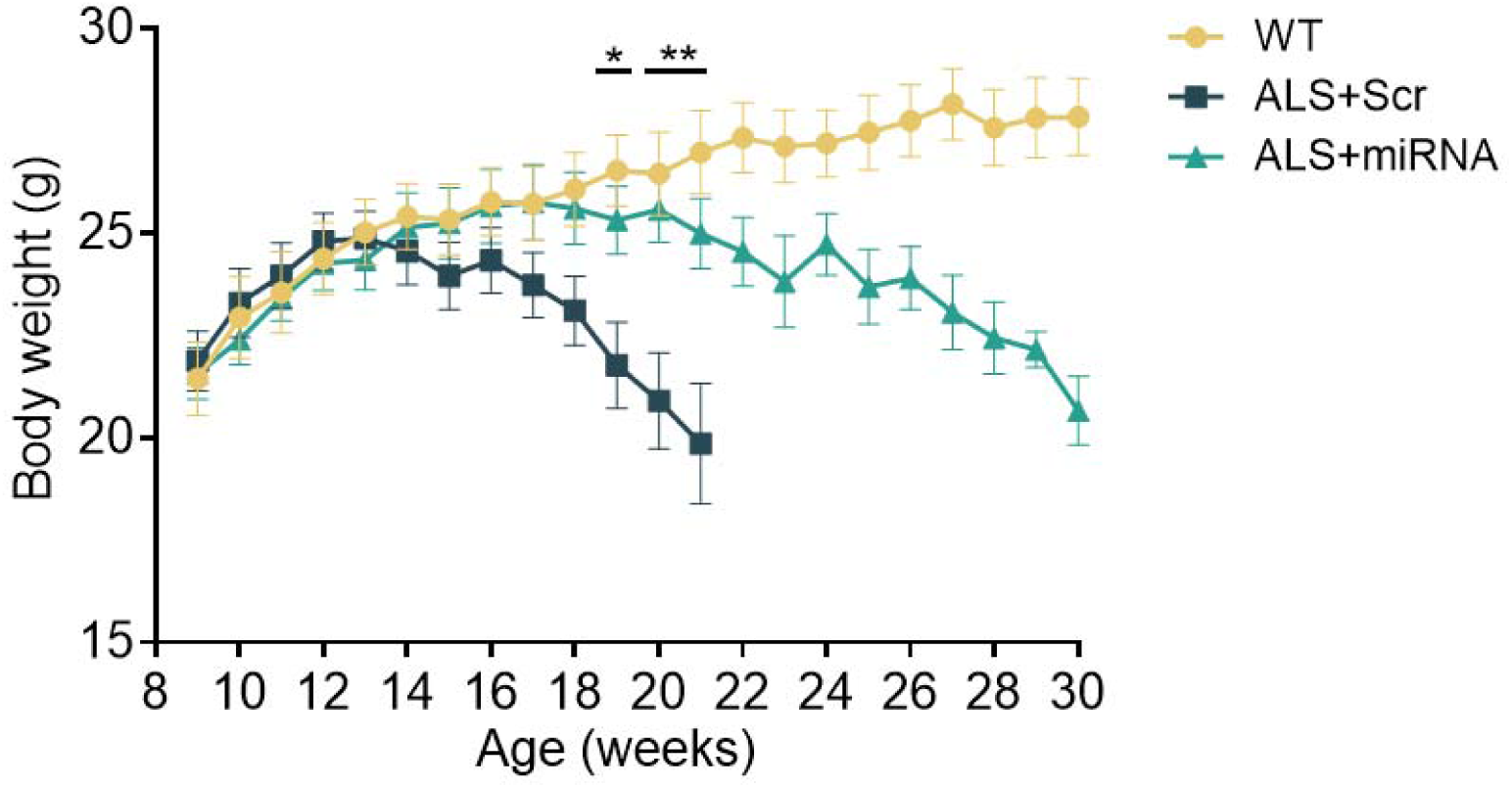
Intramuscular delivery of rAAV2-retro ameliorates weight loss in *SOD1* ^G93A^ mice. Comparison of the body weight curves of WT, rAAV2-retro-Scramble-treated, and rAAV2-retro-miRNA-treated ALS mice. n=12 mice per group; One-way ANOVA with Tukey’s post hoc test. \**P* < 0.05, \*\**P* < 0.01 for the comparison between rAAV2-retro-Scramble-treated and rAAV2-retro-miRNA-treated groups.

**Figure S3:**
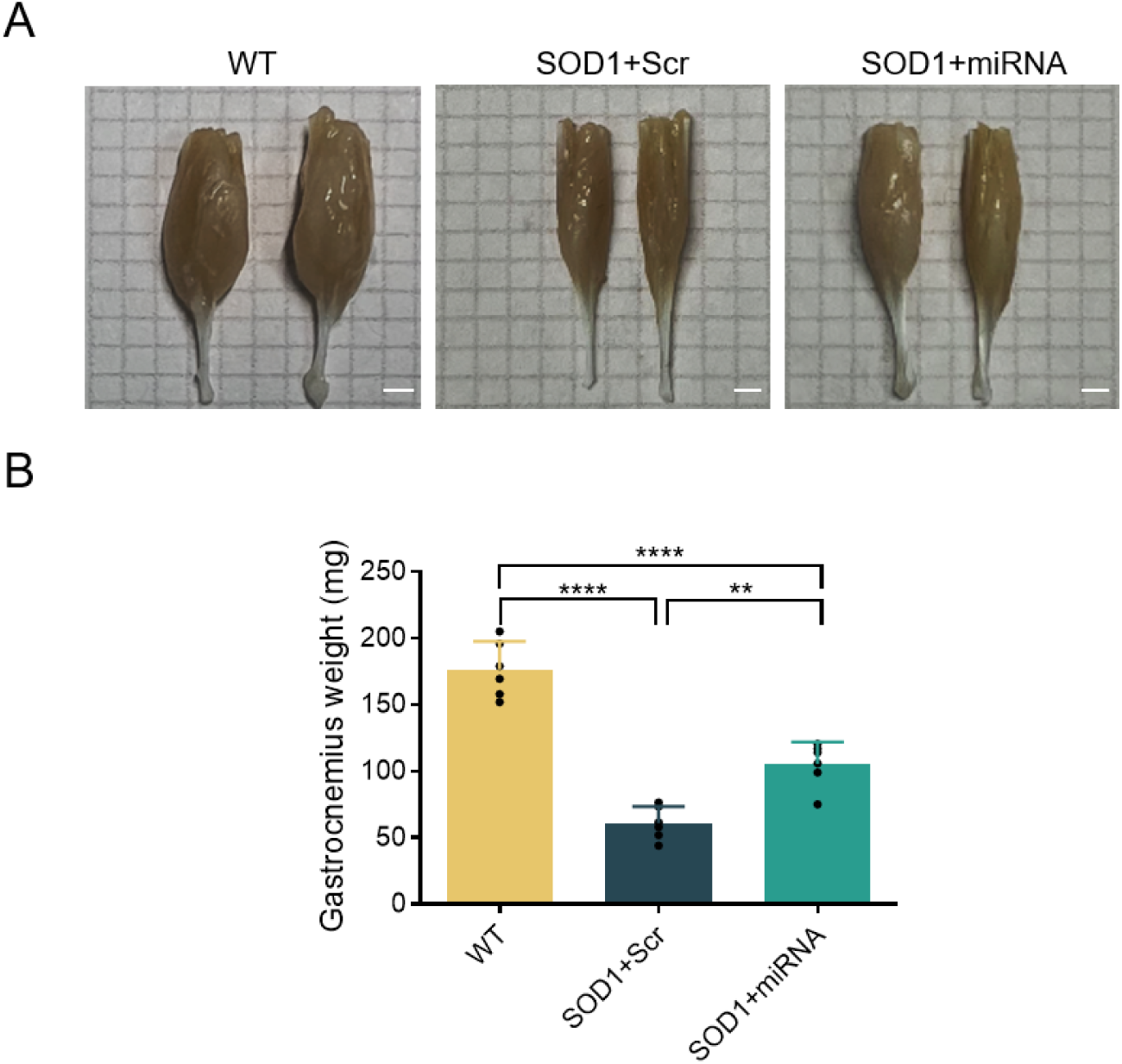
Representative images and quantitative analysis of the gastrocnemius muscle obtained at day 130 from WT and *SOD1*^G93A^ mice. (A) Gastrocnemius muscle sections from P130 WT, rAAV2-retro-Scramble-treated, and rAAV2-retro-miRNA-treated ALS mice. Scale bar, 2mm. (B) Muscle weight quantification. Data represent mean ± SEM; n = 6 mice per group. **P< 0.01, ****P < 0.0001; One-way ANOVA with Tukey’s post hoc test.

